# The ECOLOPES PLANT MODEL : a high-resolution model to simulate plant community dynamics in cities and other human-dominated and managed environments

**DOI:** 10.1101/2024.09.23.614561

**Authors:** Jens Joschinski, Isabelle Boulangeat, Mariasole Calbi, Thomas E. Hauck, Verena Vogler, Anne Mimet

## Abstract

Cities and urbanized areas have become an important habitat for organisms. Although nature can provide multiple ecosystem services to humans, urban planning and architecture often do not adequately consider the ecological needs of species. In ECOLOPES we envision buildings, and in particular building envelopes, to be a joint habitat for humans, plants, animals and microbiota as equally relevant stakeholders. A centerpiece of the planning and design tools that ECOLOPES provides is a joint soil-plant-animal community model. We present the first prototype of the ECOLOPES PLANT (community) MODEL that responds to 3-dimensional (building) geometry and management, show potential applications as a stand-alone model, and discuss its technical and conceptual limitations that are yet to be overcome.

## Introduction

Biodiversity provides multiple tangible and intangible benefits to society, including e.g. psychological (Fuller et al., 2007) and medical (Erwin et al., 2010) remedies, agricultural benefits (Dainese et al., 2019) and sources of economic wealth (Echeverri et al., 2022). The largest ecological benefit of biodiversity is probably the sustenance of ecosystem functioning, and of its temporal stability (e.g., Cardinale et al., 2012; Tilman & Downing, 1994). Overall, a sustained biodiversity loss (mass extinction) would have far-reaching consequences, including on the ecology and long-term evolution of surviving taxa (Jablonski, 2001).

Biodiversity is indeed rapidly declining globally (Díaz et al., 2019; Hallmann et al., 2017; Pereira et al., 2010; van Klink et al., 2024), at a rate comparable to historic mass extinctions (Ceballos et al., 2015) and thus of major concern for society (United Nations, 2015, SDG 14 & 15). Patterns and drivers of biodiversity loss differ among taxa (Sánchez-Bayo & Wyckhuys, 2019), but land use and land cover change is globally the largest cause of biodiversity decline (IPBES, 2019; Jaureguiberry et al., 2022). Humans are responsible for the changes in land use and land cover with highest impact on biodiversity, most notably agricultural intensification (Egli et al., 2018; Kehoe et al., 2017) and urbanization (McDonald et al., 2020; Seto et al., 2012). Stopping or even reverting the trends in land use and land cover change is a major societal challenge on its own (United Nations, 2015, SDG 2 & 11), and solutions that revert the trend would benefit not only biodiversity but also food security and human health and well-being.

The impacts of land use and land cover changes can be moderated by thoughtful interventions (e.g., hedges for connectivity, crop diversity, wildflower strips) that can significantly increase ecological stability and functioning without large negative economic impacts (Bommarco et al., 2013). Similarly, simple and cost-efficient measures (e.g. reduced management) can improve biodiversity and ecological functioning in heavily urbanized areas (Threlfall et al., 2017; Vega & Küffer, 2021). Yet, better results may be achieved when ecology is integrated and made an explicit objective, both in agriculture (organic farming) and in architecture and urban planning (e.g., regenerative design, net-positive design, biophilic design), as early consideration allows mediating conflicting objectives (Birkeland, 2020) and considering the multiscalar nature of ecological processes (Beninde et al., 2015).

The ECOLOPES (ECOlogical building enveLOPES) project aims to integrate ecology as early as possible into the design process, with the aim to design ecologically sound buildings (Weisser et al., 2023). In contrast to existing approaches, ECOLOPES considers not only the ecosystem services that urban green provides to humans, but also the perils and chances that architecture delivers to plants, animals and the microbiota. As such, it represents a radical change of view, from an anthropocentric view towards a vision in which non-human stakeholders are equally considered. Consequently, the ideal *ecolope* does not resemble a garden or park, but is a self-managed and functional ecosystem that requires no or only very little management. Moreover, ECOLOPES does not only conceptually consider ecological needs, but also provides the tools for a technical, integrated analysis that can be put in practice by landscape planners and architects (https://gitlab.com/ecolopes-team). One of the key outcomes of ECOLOPES will be a coupled ecological model, which jointly simulates plant, animal and (soil) microbe communities, their interaction, and how the community (and its temporal change) is affected by architectural features and urban planning decisions. This ecological model can in turn be combined with other computational tools to become the centerpiece of a data-driven design recommendation system, and help lowering the conflict between needs of humans and of other stakeholders.

We here present the first prototype of the ECOLOPES PLANT MODEL, as one of the parts of the envisioned joint ecological model. This C++ – Model is derived from the model Fate-HD (Boulangeat et al., 2014) and simulates plant communities in a 3-dimensional environment. We strictly limit ourselves to a technical description and discussion of capabilities of the model itself, as the integration into a complex computational workflow (plugin for Rhino®, McNeel Europe) is not finished yet. A discussion of the model (or any successors) in an applied context will accordingly be published separately.

## Methods

The ECOLOPES PLANT MODEL is a stand-alone tool that models plant dynamics at high resolution and which will also be integrated into the computational workflow of ECOLOPES. We here summarize the model by describing its modeling objectives, concepts and required inputs. We also demonstrate the model’s general utility with the help of (manually created) 2D and 3D inputs and a visualization in R. The full description of the model can be found in the appendix. This description follows the ODD (Overview, Design concepts, Details) protocol for describing individual– and agent-based models (Grimm et al., 2006, as updated by Grimm et al., 2020).

## Objectives and novelty

The ECOLOPES PLANT MODEL aims to predict the presence, abundance and community dynamics of different types of plants, usually clustered into Plant Functional Groups (PFGs). It is derived from the Fate-HD community model (Boulangeat et al., 2014). In short, Fate-HD is a landscape model that combines species distribution modeling with coarse-grained (generalized) process-based modeling, thus achieving a balance between general applicability and accuracy. The outputs of Fate-HD are not to be taken at face value, but indicate relative shifts in community composition over time.

The ECOLOPES PLANT MODEL inherits most of the concepts of Fate-HD, but repurposes it to evaluate the effect of microenvironmental conditions on community composition and succession, in a 3-dimensional environment. Like its predecessor, the current version captures relative community shifts, but in contrast to Fate-HD the long-term aim is a more accurate prediction of plant biomass in absolute terms. Moreover, the ECOLOPES PLANT MODEL is meant to be combined with other tools, e.g., an animal model that predicts animal community dynamics based on the plant community, and hence requires a high degree of interoperability and flexibility. This version of the model serves as backbone for future refined versions and identifies technical constraints for providing more accurate results. To this aim, we revised the technical implementation while conserving almost all concepts from the precursor, except for light competition, which is further elaborated in the discussion.

In summary, this version of the model is not meant to provide information about biomass distribution in absolute numbers yet, but it allows showcasing the technical feasibility and utility of a 3D community model, and it allows identifying conceptual challenges that need to be solved in the future.

## Entities and scale

The model is designed for use in urban environments, and it simulates community dynamics over the typical lifespan of a building envelope; its spatial resolution is fixed at 1m x 1m x 1m, while the extent is variable but shall be in the order of 100 x 100 x 10 m (length, width, elevation). It is made up of cubic cells in which a plant community can thrive. The cubic cell itself is stratified, mimicking the stratification of plant communities. We chose four strata in our tests, corresponding to herb, shrub, understory and canopy layers. Plants differ in maximum sizes and growth forms (e.g. grasses vs. shrubs vs. trees), and hence both in the number of strata they grow through and in the time they stay within each layer.

The principal agent of the model is the demographic composition of a local stand of plants of the same species of functional group (i.e. not explicitly an individual plant). In lack of a better wording, we will name this local aggregation of plants of the same taxonomic group but different ages a deme (but without implying any genetic structure or local adaptation). The deme itself consists of age cohorts of individuals, but individuals within an age cohort are not uniquely identifiable (no inter-individual variation). Multiple demes of different functional groups or taxonomic identity can potentially live together in a cell to create a community, and each is defined by its own trait values (e.g., demographic traits’ values, responses to disturbances, etc; see Supp. S1, Table 4). The trait values have to be provided by the user, and one way to overcome data limitations is the aggregation of functionally redundant species into the same group, i.e., the creation of Plant Functional Groups (Boulangeat et al., 2012).

Plant composition dynamics is implemented through the temporal simulation of demographical changes of the plant community (i.e., temporal changes of deme abundances) in each stratum of the cells, also accounting for the simulated concomitant changes in light availability (caused by changes in deme abundance). We use a slightly different and more explicit concept of light in this model than the precursor Fate-HD, and we will elaborate the differences in the discussion. Briefly put, light passes through the strata of the community and gets reduced along the way. In contrast to Fate-HD, where it is not available light that is in the reasoning but rather shading effect through the evolution of the canopy, in this model the amount of available light is corrected by shading (e.g., by buildings). This ensures that shaded cells fall earlier to “medium” or “low” light conditions and potentially exclude non-tolerant

PFGs. Furthermore, light may fall at an angle below 90°. If this is the case, it will partially pass through the neighboring cell and is accordingly reduced by the neighbor’s plant abundance. Apart from the dynamically changing light conditions, the plant demes also respond to the following microenvironmental (cell) parameters that are usually provided as an input to the model, adapted from the “habitat suitability” and “disturbance” modules of FATE-HD:

– *Soil depth*. The depth of the soil is a limiting factor for plants, especially deep-rooting ones. Accordingly, plants whose rooting depth exceeds the soil depth are unable to grow.
– *Soil class*. Plants differ in soil requirements and hence have different soil profiles. If the soil class in a cell does not match the plant’s profile, the plant is unable to germinate or grow.
– *Disturbance*. External, user-defined disturbances such as herbivory, fire or management can impact population demography. The strength of the effect may differ among PFGs.
– *Shading*. Shading caused by buildings or natural structures (cliffs) reduces available light and thereby intensifies competition among PFGs (as explained above).

These four cell attributes are constant throughout the model run in the stand-alone version of the new model. However, the model may also be compiled as a shared library (.dll), allowing it to be embedded in other programs and models (not available under Windows). If used in such a way, the inputs “soil class” and “disturbance” can be exchanged at every time step. This allows, e.g., modeling directional and stochastic climate change through changes in the soil class attribute.

## Process overview

The model largely works on individual cells (Fig. 1) and uses the process employed by Fate-HD: in each time step, each plant deme is disturbed by a number of fixed, annual disturbances. Depending on its PFG traits, each growth stage of a deme may react differently to the disturbances, including not being affected at all. Then it is checked, separately for each stratum, whether the current light conditions are sufficient. A common outcome is the death of all lower plants, while the upper plants survive. Subsequently the deme ages by one time step, potentially causing the oldest demographic group to die. Aging is followed by a calculation of the new light conditions. Light is calculated based on the demography of the whole community of the cell, and in contrast to FATE-HD, also of the neighboring cell if enabled. This is actually more important for the targeted resolution (1m), as individuals could spread over several cells. Germination and recruitment are then individually computed for each deme, and the outcome is a number of newly produced seeds. These processes depend on the current suitability of the cell, which is in turn determined by soil depth and soil class. The newly dispersed seeds are not immediately placed back into the cell. Instead, all seeds across the landscape are collected and then uniformly and randomly dispersed across the site. The Fate-HD modules “soil”, “drought”, “fire” and “aliens” of Fate-HD were not ported to this version, but can be emulated by other means; “disturbance” was reduced to removal of adult plant material (no seed predation and no resprouting); and “dispersal” was simplified to accommodate the smaller spatial scale. See appendix S1 for details.

**Fig. 1:**
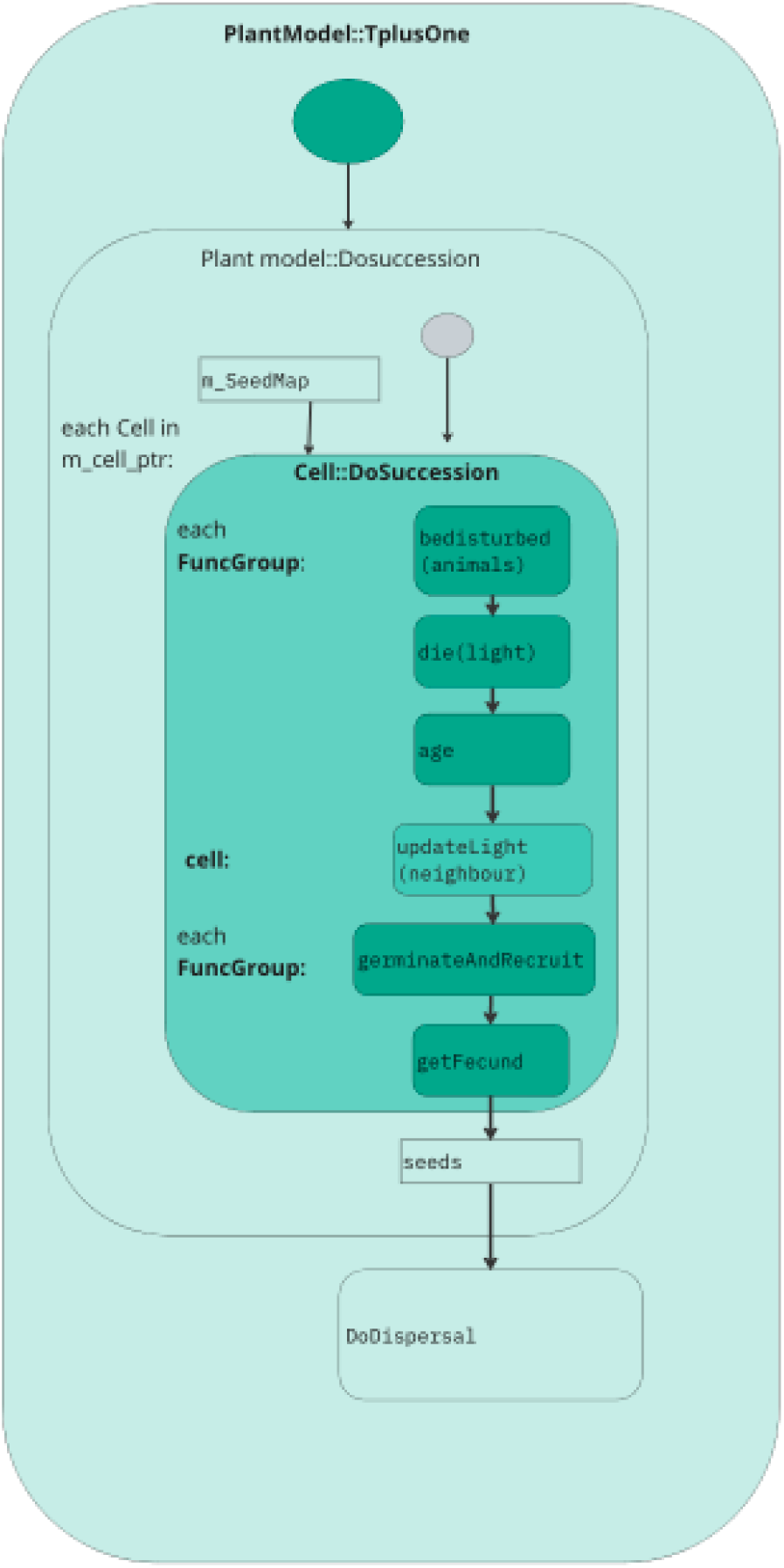
Process overview. The order of events is largely copied from Fate-HD, so that conceptual changes to the model are kept to a minimum. Processes in this diagram match function names of code (see UML diagram in Supp. S1); the deme is called FuncGroup in the code.

## Input and output data, initialization

The model requires the following information to run:

– Configuration parameters (12 variables, see Supp. S1 Table 3)
– Definition of Plant Functional Groups, including their response to disturbances (e.g. management, animals) and soil (26 variables, see Supp. S1 Table 4)
– Microenvironmental information for each cell. This currently includes the amount of shading, soil class and soil depth, and the amount of each (user-defined) disturbance.

The output of the model is an abundance value of each deme in each cell, for each time step that was specified. The abundance values are unitless expressions to compare among PFGs, but in the future are expected to correlate with plant biomasses (see discussion).

Special consideration needs to be given to sensible starting populations. Starting conditions may include, e.g., a barren soil that gets colonized over time; a random starting population; an initialization period of 1000 time steps on either of above, to achieve a stable community; the use of ecological mapping to explicitly instantiate each plant with correct age; or a combination of above. Longer initialization periods are in principle possible, but increase the runtime of the model, while explicit instantiation requires further high-resolution data. We thus decided to initialize the models with zero – and 1-year old plants from each PFG, with an abundance of 1 to 100 (uniformly random) wherever the habitat is suitable. The model then runs for five time steps to achieve more variation in the age distribution in each cell. We are aware that the required inputs (shading, depth, soil classes, PFGs and configuration parameters) are difficult to create manually, and that the manual use of the tool is tedious. We provide a short user guide in the supplementary material (Supp. S2) to explain the required inputs, and steps to perform and run the model, but we would like to emphasize that separate tools and workflows are in preparation (led by Verena Vogler), which will considerably simplify input creation and visualization of results in the future.

## Model testing

We use the gtest framework (<u>google.github.io/googletest/</u>) for unit and integration tests, but tests are currently limited to the most crucial functions (40 tests in 6 test suites), focusing in particular on the completely revised higher-level classes. Although we also revised the model architecture and class relationships, low-level functions were changed only very sparingly. We assume that these low-level functions were already rigorously tested in the predecessor model (Fate-HD) and performed only informal tests. Where applicable, we provide suggestions for future revision (see “issues” in repository, tag “coding issues”).

The most informative test suite regards a model run on a landscape of 18 cells, covering management, habitat suitability, and disturbance, but not soil depth; soil depth was, however, tested informally and is known to produce valid results. We also disabled dispersal, because it randomizes results and is inherently more difficult to test. Test results can be verified by inspecting the file “PlantModelTestNoDisp.cpp” (folder “tests”) on the repository, or by running the model with the file “test_noDisp.json” and comparing inputs and outputs manually.

## Data preparation and visualization

To demonstrate the model’s utility but also to emphasize its current limitations, we prepared two examples. For both examples we created 10 Plant Functional Groups (PFGs) defining the traits of the plants. We used 10 randomly drawn PFGs from a dataset that creates generic PFGs with worldwide applicability (Calbi et al., 2024), but adapted and simplified them to our use case. The PFGs differ in soil type and depth requirements and in parameters relating to growth, but not in shade tolerance (table 1).

**Table 1.**
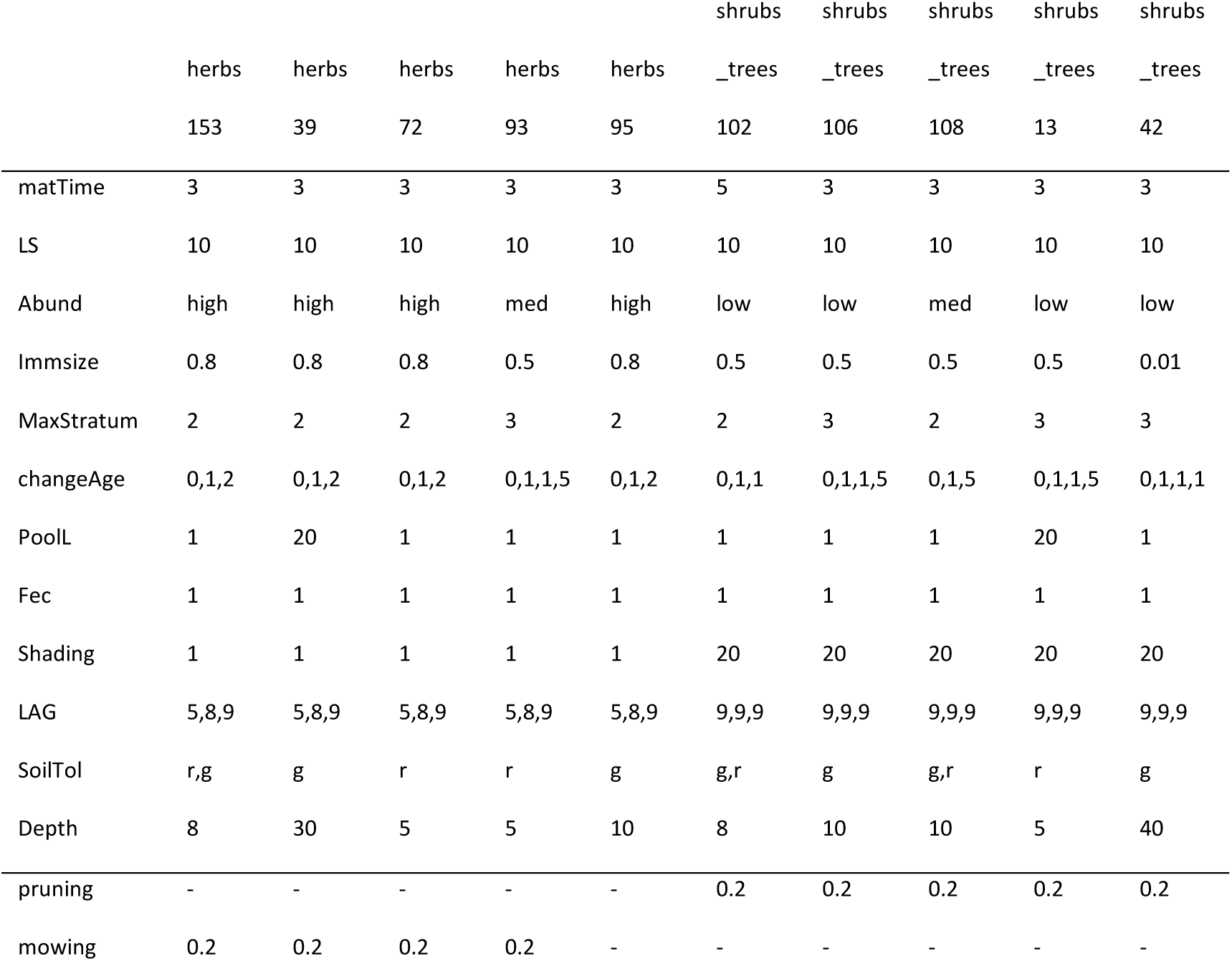
Attributes of the 10 example PFGs. “Pruning” and “mowing” represent effect sizes of management decision (annual removal rates), for other parameters see supplementary material (Supp. S1). Light tolerance was the same for all PFGs (no tolerance for shade in any PFG, except in germinant stage).

In the first example we use a 2-dimensional planar surface without shade. We defined two management plans, called “mowing” and “tree pruning”, and applied them annually in the western and in the eastern side of the site, respectively. Mowing affects the herbs, while shrub/tree PFGs are affected by pruning. PFG “herbs_95” is not affected by either management plan. In the other direction (north to south), we apply a gradient of soil depth from 0 to 45 cm in steps of 5 cm.

The second example is a 3-dimensional input geometry: a cuboid with a size of 10 x 10 x 5 m lies on a planar surface of 20 x 20 m. Light immediately north of the cuboid is reduced due to shading (10 – 50% reduction). Two balconies are attached to the southern face of the cuboid, creating also a low amount of shading underneath, and mimicking a more complex topography. The soil has a depth of 50 cm on the ground and on the balconies, and a depth of 10 cm on the top of the cuboid; tiles along the walls of the cuboid do not contain any soil. We further define two soil types, which we term “ground” and “roof”, and which are distributed on the ground and on the cuboid, respectively. We provide an intermediate amount of general management (0.5) everywhere.

The outputs are visualized using R version 4.4.0, “Puppy Cup”(R Core Team, 2024) and the packages jsonlite (Ooms, 2014), tidyverse (Wickham et al., 2019) and rgl (Murdoch et al., 2024). The R script for the entire procedure is available in the repository and in Supp. S3.

## Results

All models since version 0.5.1 pass all unit and integration tests. This means that the model is fully functional and that the tested sections work within the specifications set out in the introduction and the ODD (see Supp. S1): the model creates a temporally changing and spatially variable plant community, which is affected by soil depth, soil class, shading conditions and disturbances (e.g. management decisions or herbivory); the effect of management on the communities can be calculated manually based on the inputs (as is done in the unit tests), while soil depth and soil class produce a binary response (match / mismatch).

The effect of shading on competition among PFG is non-linear and too complex to be included in simple unit tests but can be verified by manual inspection of the results (see below). In the first (2-dimensional) use case we observed a slight increase in total abundance (summed over all PFGs) and soil depth (data not shown). The effect was, however, strongly contingent on the PFGs that were used, and individual PFGs differed in patterns. For instance, PFG herbs 39” only occurred on soils with a depth of 30 cm or more but did otherwise not vary with soil depth (Fig. 2A, left). PFG “shrubs_trees 106” occurred from 10 cm onwards, but its abundance decreased slightly with depth (Fig. 2A, right). Both groups of PFGs were also affected by management, in particular by tree pruning, with differences in the direction of the effect (Fig. 2B). There was no consistent change of abundances or community dynamics over time, except that abundances were reduced over the first two time steps. There was strong cyclic turnover in the community composition, however (data not shown). The period of the cycles was two for all PFGs, so PFG abundances were at all times either in synchrony or exact opposition. Cycles of adjacent cells were not coupled, unless light was allowed to pass at low angles through neighboring cells – in this case, regular patterns occurred in the spatial distribution of plants.

**Fig. 2:**
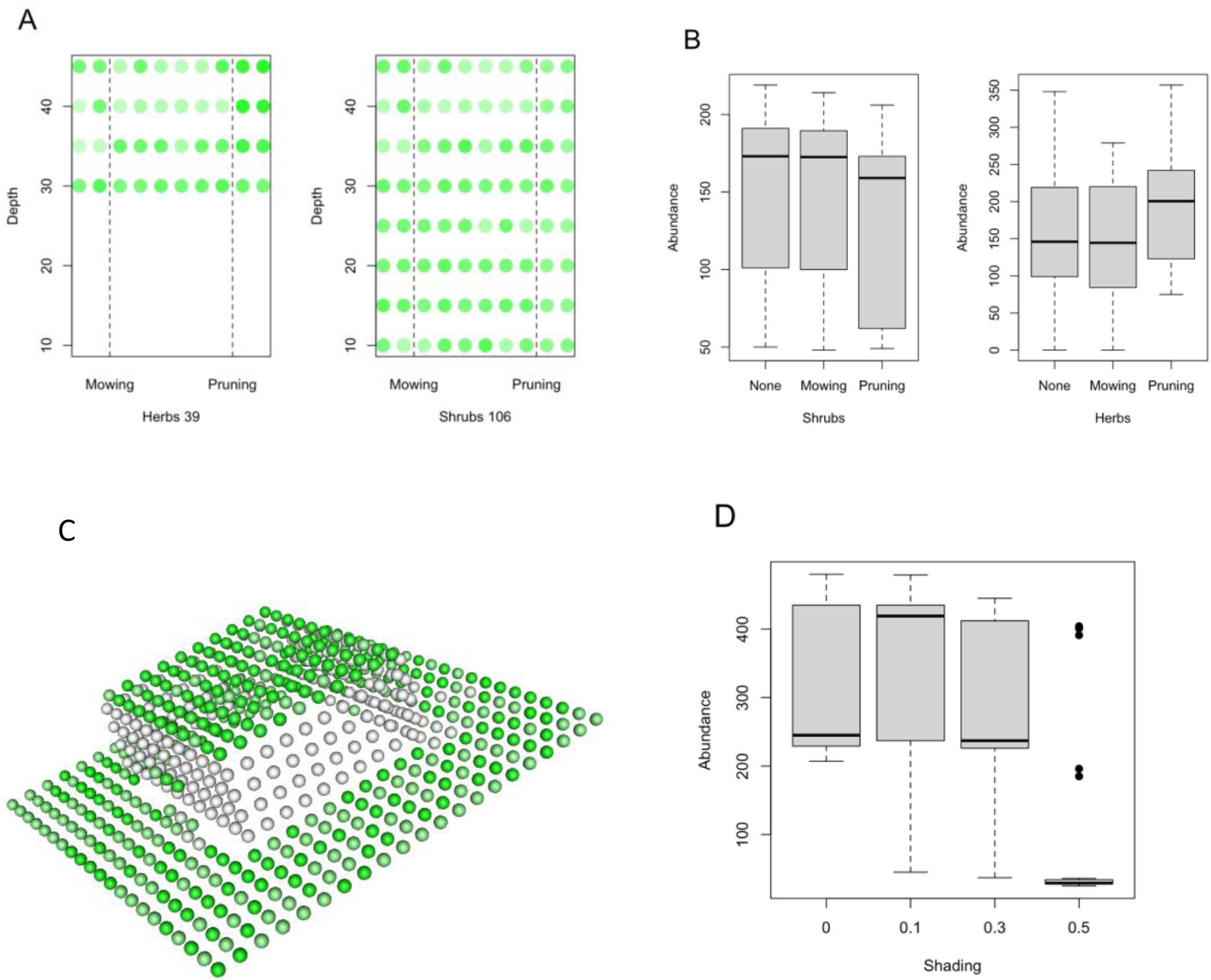
Graphical representation of modeling results. A) 2-dimensional planar surface without shade and with soil depth gradient from north to south. Mowing occurs on the western side, tree pruning on the eastern side. Color intensity indicates total abundance of the two PFGs “herbs 39” and “shrubs_trees 106”, summed over time B) Comparison of summed PFGs abundance between the three management types of A), separately for all shrubs and all herbs. C) 3-Dimensional input geometry. Shading is present immediately north of the building, soil depth and soil classes vary between roof and ground level, and general management is present on the site (see methods for full description). For demonstration purposes, summed abundance across all PFGs at time step 5 is shown. D) boxplot of summed abundance against amount of shading for C).

Using the second example topography presented in the methods, the spatial (3D) configuration of the community mirrored the input topography, and the effects of light and soil are readily visible in the outputs (Fig. 2C, D). For instance, shading (north of the cuboid) caused lower total abundances (Fig. 2C) and herbal plants were disproportionately reduced (herbs contributed on average 53.9% of the cell’s PFGs abundance under full light, 12.2% in shade; data not shown); further, all PFGs were missing along the walls of the cuboid, because the soil class did not match with the PFG requirements. Overall, higher amounts of shading had a strong effect on total PFGs abundance (Fig. 2D).

## Discussion

Before discussing the constraints, limitations and broader relevance of the model, the first question should be whether the model actually behaves as intended and conforms to the specified goals and concepts.

The ECOLOPES PLANT MODEL is part of a complex computational toolchain, and while rigorous software engineering principles are useful for nearly any individual-based model, their utility exponentiates in a complex software setting with multiple components (e.g., Scheller et al., 2010; Trisovic et al., 2022). Given these requirements, and its intended use in commercially relevant applications, our model has to be held to particularly high standards regarding code conformity and stability. It does by no means reach this much higher quality threshold yet, and a provision of quality metrics (e.g. test coverage, cyclomatic complexity, static analysis outputs) would be premature. Nevertheless, we embraced critical software engineering principles wherever possible (encapsulation, clean code, assertions; full documentation and logging; version control, automated integration and testing, portability; use of standard tools, libraries and file formats) and we are confident that the software provides a reasonable basis for future development. Indeed, the unit tests ensure alignment of the code with core model specifications, and first simulation results look promising (Fig. 2). In the future more tests shall be added, including not only those unit tests which are currently missing, but also full integration tests that survey light competition outcomes in more detail. For now we suggest that the model does largely behave as intended, though a risk for unforeseen bugs remains. Unfortunately, biological research is somewhat detached from software engineering and practitioners are frequently not trained in best practices (Vedder et al., 2021), nor is there adequate consideration from funding and supervising bodies, or particular awareness in the publication process (but see Freckleton, 2018). The resulting code quality and potential error rate in biological research is disappointing (Darriba et al., 2018; Trisovic et al., 2022). On the other hand, scientists are trained in critical analysis and should be able to detect major errors and spurious results, somewhat mitigating the high risk for errors. We are confident that our model can compete against this (low) baseline of model expectations, and we carefully suggest that the model is ready for scientific use.

Having discussed the coherence of concepts and implementation, one may now ask whether the conceptual decisions themselves are sensible. The concepts were in most parts borrowed from Fate-HD and are defended in the model’s description (Boulangeat et al., 2014), but there are two important additional considerations which any reader and user of the model must be made aware of.

First, the model’s meaning of “light” differs slightly from that of Fate-HD. Fate-HD is a landscape community model, which finds a balance between complexity and generality by modeling general patterns at coarse resolution. Although both the model description (Boulangeat et al., 2014) and the code of Fate-HD mention “light conditions” and “shading” variables, they are meant to represent canopy effects, i.e., the integrated effect of shading, humidity changes, and other abiotic and biotic factors. Similarly, the output of Fate-HD (abundance) shall not be interpreted as biomass units in absolute numbers, but as a general guidance regarding relative abundances of PFGs. A future version of the ECOLOPES PLANT MODEL is, on the other hand, meant to provide accurate results in absolute values. The use of unitless abundance and canopy values hinders transition to explicit biomasses, so light and shading are taken more literally in the new model. This conceptual change has important repercussions on the interpretation of results. For instance, in Fate-HD it was perfectly logical if an understory plant required a canopy for survival (facilitation), because the canopy provides a cooler and more humid climate; in the ECOLOPES PLANT MODEL it would not make biological sense if a plant could survive under shaded, but not under full-light conditions. The models hence differ in the expected parametrization.

Using the more literal interpretation of light/shading, the ECOLOPES PLANT MODEL was able to make two further additions to the concepts and implementation. First, the amount of available light is corrected by a shading index. This ensures that shaded cells fall earlier to “medium” or “low” light conditions. Secondly, light may fall at an angle below 90°. If this is the case, light willl partially pass through the neighboring cell and is accordingly reduced by the neighbor’s plant abundance. These conceptual changes to light (along with their technical implementation) ensure a more realistic treatment of shading, including shading by buildings and other geometries (Fig. 2C). Nevertheless, while the new treatment of light is an important step towards more explicit model outcomes in the future, we still advice against interpreting them in absolute terms in the current version. For instance, within ECOLOPES (Weisser et al., 2023) the model is expected to deliver «good enough» predictions on how building geometry alters community structure, such that the model can be relied upon to provide suggestions for improving building designs – but it cannot be used yet to calculate expected weight loads on green roofs.

Secondly the ECOLOPES PLANT MODEL is applied on a much smaller spatial and temporal scale than Fate-HD. In Fate-HD the community of each cell is usually a larger group of individuals, but by reducing the spatial extent to a cube meter in this model, we simulate only a low number of individual plants. This leads to a series of conceptual problems and mismatches with reality:

1) in Fate-HD the highest stratum provides a canopy that (correctly) shades the lower strata, potentially causing lower-growing plant material to die. Fate-HD does not strictly differentiate between biomass and abundance of individuals (see Supp. S1, section 9), but because the number of individuals per cell tends to be high, the lower-growing plant material of the same PFG can statistically be interpreted as being a different individual (i.e., offspring). Our model reduces the number of individuals per cell, and the abundance values must accordingly be more strongly interpreted as biomass values of the same plant. This causes an unrealistic shift from inter-to intra-individual competition;
2) Fate-HD does not model competition for space explicitly, so plants that grow within the same stratum, which do not compete for light, do not compete with each other at all; in our model the small spatial scale makes competition for space more important and should be considered;
3) Plant material that grows to a higher stratum will hover in the air and its lower strata cast no shade. This does not matter for the canopy calculation in Fate-HD (except at forest edges), but becomes important when light enters at an angle;
4) Each plant community is confined to a single cell. Plant material that outgrows a cell does not cast shade on the neighboring cell, so the results may become unreliable when the model is used for larger PFGs;
5) The highest-growing plant material will usually determine the light conditions of the cell. As long as the abundance in the highest stratum is high, nearly no plant material will grow below. Because there is no inter-individual variation in lifespan, the abundance will change suddenly, making room for other PFGs on the lowest stratum and causing the observed strong population cycling on small time scales (see results), at least when the number of PFGs and the variation in their demographics is low.

Overall, the reduction of the spatial scale warrants a revision of some of the key model concepts. We argue that the core of the problem lies in the use of a (demographic) community model for individual plants. Adding individuals with their own resource pools and (spatially explicit) biomasses should solve various issues regarding light competition and may also simplify parametrization. Moreover, a revision would solve various technical challenges regarding data storage (see Supp. S1 and issues section in the repository). A later version of an individual-based model could then add more realistic lighting calculations, including along sloped surfaces and facades.

The question how well the conceptual decisions made here reflect reality must ultimately be answered by parametrization and validation against real-world data. The parametrization is, however, cumbersome in both Fate-HD and the new model. Six important parameters are set via a configuration file (Light thresholds, abundances, fecundity; see Supp. S1, Table 3), and 14 further types of parameters (>25 entries) are specific for each PFG (Supp. S1, Table 4). While some of the PFG parameters are straight-forward to understand (e.g. lifespan), others do not have an intuitive or well-supported relationship with PFGs inherent features (size of immatures relative to adults) or are not meant to represent a measurable entity(e.g., light thresholds, a fecundity constant). Because the model parametrization by PFGs requires considerable effort (Boulangeat et al., 2012), and because there are further technical and conceptual issues to solve first (see above), we have so far opted against a separate parametrization and validation of the model. We think that the validation performed by Fate-HD shall still be mostly valid for larger spatial scales and lower resolutions, while for lower scales and higher resolutions significant work is likely still required.

Having discussed the technical implementation and the concepts, the last remaining question is the potential application of the model. As an integrated tool (Weisser et al., 2023), the model is not, in fact, immediately useful in its own right, but requires other means to generate inputs and visualize the results. Nevertheless, many researchers are proficient with R, and when combined with R scripts, one can already create meaningful inputs and examine and analyze the results. The simple example presented in the results shows that all key processes are working (Fig. 2). First, PFGs differ in the spatial distribution along the site, according to soil class and depth tolerances (Fig. 2A). Secondly, specific plot regions can be affected by disturbances, and the effect of the disturbance is PFG-specific (Fig. 2B). Thirdly, the three-dimensional setup of the site is adequately considered (Fig. 2C). For instance, total plant biomass is reduced in shade (Fig 2D). The plant communities growing in the shade underneath the balcony also differ from those growing above them, showing the potential to include complex topographies. In summary, the example demonstrates that meaningful inputs can be created manually, and visualized or analyzed with R.

The example presented here was on purpose very simple, as it requires the manual input of data. Yet, the workflow could also be used for more complex use cases, for example to model plant communities in alpine or otherwise topographically complex habitats. A digital elevation model (DEM), analyzed with a shading function (e.g. hillShade, package raster) should provide shading values, and combined with maps of soil depth and soil classes it should be able to provide all necessary inputs for the model. Further, site-specific abiotic and biotic stressors (e.g., strong winds, fire, herbivory pressure) could be added via custom-made disturbance maps. The process is admittedly quite labor-intensive though and requires site-specific scripting and tweaking of the data.

## Conclusion

The current conceptual limitations notwithstanding, the model is technically ready and can already provide a reasonable plant community that reacts to microenvironment and topography. A tool built on this model or any successor that solves the presented issues (Vogler, Grasshopper plugin in Rhino) can help visualizing the effect of the urban fabric on plant communities, and especially with future improvements in mind, will help develop buildings that promote ecological functions (Weisser et al., 2023).

## Author contributions

AM and TH conceived the original idea and led the project. IB provided the orginal model (FATE-HD) and contributed to the conceptual framework. VV provided help in data structuring, inputs and outputs, and performed visual validations of earlier model outputs. MC provided additional conceptual support, in particular regarding model parametrization.

## Acknowledgments

We gratefully acknowledge funding by the EU H2020 FET-OPEN project ECOLOPES (Grant agreement number 964414). We further thank Laura Windorfer, Victoria Culshaw, Wolfgang Weisser and the whole ECOLOPES consortium for helpful discusssions regarding the conceptual development.

