## Supplementary material for "The ECOLOPES PLANT MODEL : a high-resolution model to simulate plant community dynamics in cities and other human-dominated and managed environments": Supp. S1

### The ECOLOPES Plant Model - ODD

Jens Joschinski, Studio Animal-Aided Design

This report has been written by Jens Joschinski. Some text is copied or adapted from (non-public or public) documents of the ECOLOPES project. ECOLOPES has granted permission to use the text presented herein.

Copyright (C) 2022 - present Studio Animal-Aided Design
Permission is granted to copy, distribute and/or modify this document under the terms of the GNU Free Documentation License, Version 1.3 or any later version published by the Free Software Foundation; with no Invariant Sections, no Front-Cover Texts, and no Back-Cover Texts. A copy of the license is included in the section entitled "GNU Free Documentation License".

#### Preface

This model description loosely follows the ODD (Overview, Design concepts, Details) protocol for describing individual- and agent-based models (Grimm et al., 2006), as updated by Grimm et al. (2020). It concerns the ECOLOPES PLANT MODEL, version 1.0.2, which can be found at https://gitlab.com/ecolopes-team/plantmodel. Documentation for older versions of the same model can be found by browsing the commit history and checking the respective README files. The repository also contains an "issues" section, which reports bugs, missing tests or other coding issues, but also conceptual limitations and oversights. This documentation refers to the issues where applicable.

#### Context

This model is developed as part of the ECOLOPES ECOLOGICAL MODEL, which is used for designing building envelopes with ecological function (WEISSER ET AL., 2023). The ECOLOPES ECOLOGICAL MODEL is a spatial-explicit model that models the interdependent spatial and temporal dynamics of the soil, microbiota, plants, and animals, as response to the regional species pool, the local abiotic conditions, the geometry of the building, the substrate used to design the ecolope, and the management. The biological units of the model are plant (PFG) and animal (AFG) functional groups, i.e. species with similar ecological function, and generic soil classes, which makes the model generalizable to all sets of conditions. The model is based on a multiscalar approach: the regional model determines which FGs of the species pool have a reasonable chance to colonize the ecolope according to its location in the city; the local model applies a second filter on these species based on the abiotic and biotic conditions delivered by the ecolope, and simulates demographic processes to predict the outcome of inter-and intra-specific interactions. The output of the models is a temporal sequence of plant-animal-soil community development. The ECOLOPES models and data can be found at https://gitlab.com/ecolopes-team and https://gitlab.com/ECOLOPES. Although the ECOLOPES PLANT MODEL is part of the larger ECOLOPES ECOLOGICAL MODEL, it could (in principle) also be used as a stand-alone tool, or repurposed for other research needs. This document only describes the Plant model.

### 1 Purpose and Patterns

The ECOLOPES plant model primarily aims to predict the presence, abundance and community dynamics of different types of plants, usually clustered into Plant Functional Groups (PFGs). It is derived from the Fate-HD community model (Boulangeat et al., 2014), which is a landscape model that includes spatiotemporal dynamics; considers multiple species or species groups in interactions; and accounts for the processes shaping biodiversity distribution. By combining species distribution modelling with coarse-grained (generalized) process-based modelling, Fate-HD achieves a balance between general applicability and accuracy. The outputs are accordingly not to be taken at face value, but indicate relative shifts in community composition over time.

The ECOLOPES plant model inherits most of the concepts of Fate-HD, but repurposes it to evaluate the effect of microenvironmental conditions on community composition and succession, in a 3-dimensional environment. It is meant to be combined with other tools and hence requires a high degree of interoperability and flexibility. In particular, the plant model:

1. Is able to receive inputs from, and provide outputs for the 3D CAD software Rhino®. McNeel plans to release the model (or a successor) as free plug-in via Food4Rhino.
2. Is able to communicate with separate animal and soil models. The three models together form the ECOLOPES ECOLOGICAL MODEL.
3. is parametrized with general-purpose Plant Functional Groups.

Like its predecessor, the current version captures relative community shifts, but in contrast to Fate-HD the long-term aim is a more accurate prediction of plant biomass in absolute terms. This version of the model serves as technical backbone for future refined versions. So, while this version of the model cannot provide information about biomass distribution in absolute numbers yet, it allows showcasing the technical feasibility and utility of the approach, and it allows identifying conceptual challenges that need to be solved in the future.

### 2 Entities, state variables, and scales

#### 2.1 Spatial and temporal resolution

FATE-HD did not constrain spatial extent, temporal extent, or spatial resolution in any way, but was designed as a landscape scale model with annual time scale. The ECOLOPES PLANT MODEL, on the other hand, is tuned for use in urban environments (while still being applicable to any other ecosystem), and it simulates community dynamics over the typical life span of a building envelope. I assume that the simulation time ranges from 10 to 200 years, the spatial extent is in the order of 100 x 100 x 10 cells, and the resolution of cells is 1 $m^2$ (Table 1).

Table 1: Spatial and temporal dimensions.

| **Dimension** | **Value range** |
|---|---|
| Spatial extent | 50 x 50 x 1 - 200 x 200 x 10 cells |
| Temporal extent | 10-200 time steps |
| Spatial resolution | 1m x 1m x 1m |
| Temporal resolution | 1 year |

The model simulates 1 x 1 m cells in a 3D landscape. Not all cells need to be present (e.g. building interior, air between buildings), and undercuts or balconies can also be modelled. The maximum extent in x-, y- and z- direction is user-defined. Each cell is a cube in which a plant community can thrive. The cell itself is stratified into four layers, though the number of strata can also be modified. Plants differ in maximum sizes and growth patterns and hence both in the number of layers they grow through and in the time they stay within each layer. For example, grasses may stay in the lowest stratum throughout their life span; shrubs reach the second or third stratum quickly but do not grow further; and trees may reach the highest stratum but take longer for their growth through the strata.
The model contains no checks that strata sum to 1 m height, and higher cells would collude with the cell above. However, any 3D dynamics (competition for light) are expected to happen within the 1 $m^3$ cell, among cells of the same horizontal plane, or among all cells but without considering space (random dispersal of plants), so there generally is no interaction among stacked cells. Thus, plants growing larger than 1 m can be easily accommodated, as long as there is no second layer of voxels within reach above them (undercuts or balconies).

Despite the expectations mentioned above, one can nevertheless use the model outside urban context, and one can change both the simulation duration and the spatial extent; however, the assumption that all seeds disperse freely across the site becomes unrealistic if working on a too large spatial extent.

The model does not incorporate the resolution explicitly into any calculations (except for an experimental calculation of light falling through neighboring cells), but a length and width that is much smaller than the size of one plant results in awkward outcomes, because the model does not provision for the case that plants grow out of their cell. The model does not incorporate seasonality and does not include fine-scaled modelling of growth processes within a year, so a temporal resolution below 1 year is not yet possible.

#### 2.2 Entities

Table 2: Entities in the model

| Entity | Description |
|---|---|
| Environment | (urban) habitat containing cells, global simulation parameters and regional model information |
| Cell | a 1 x 1 x 1 m cell with environmental information and plant demes |
| Plant deme (FuncGroup) | Simulates the demography dynamics of a PFG |
| Plant Functional Group (PFG) attributes | Defines habitat suitability, growth, competition and disturbance attributes. Provided by the user. |

##### Individuals, functional groups and populations

The principal agent of the model is not explicitly an individual plant but the demographic composition of a local stand of plants of the same functional group. In lack of a better wording, we will name this local aggregation of plants of the same taxonomic group but different ages a deme (but without implying any genetic structure or local adaptation), called FuncGroup in the code. Multiple such demes can potentially live together in a cell and create a community, and each is defined by its own demographic traits, responses to disturbances, etc. The deme itself consists of age cohorts of individuals, but all model processes act on the cohort size; in other words, there is no inter-individual variation or possibility to target specific individuals. The traits have to be provided by the user, and one way to overcome data limitations is the aggregation of functionally redundant species into the same group (Boulangeat et al., 2012). In accordance with FATE-HD, the entity providing plant attributes is termed Plant Functional Group (PFG) throughout the code and this documentation; nevertheless, no assumptions are made regarding the creation of the data, and it would also be possible to use the model with species instead of PFGs.

For clarity we use the words “deme” and “cohort” to indicate modelled entities, “PFG” for the data that is used to model demes, and “plant” for the real-world entities that we attempt to simulate.

##### Cells, microenvironment and landscape processes

The demes live in stratified cells, and grow according to their demographic characteristics and environmental influences. The following environmental parameters are stored in each cell:

- *Soil depth*. The depth of the soil can be a limiting factor for deep-rooting plants. Accordingly, PFGs whose rooting depth exceeds the soil depth are unable to grow.
- *Soil class*. Plants differ in soil requirements and hence have different soil profiles. If the soil class in a cell does not match the PFG’s profile, it is unable to germinate or grow. The soil class is considered to be homogenous throughout the cell’s depth, i.e., there is no stratification of soils.

- *Shading*. Shading caused by buildings or natural structures (cliffs) reduces available light and thereby intensifies competition among PFGs.
- *Disturbance*. External, user-defined disturbances such as herbivory, fire or management can impact population demography. The strength of the effect may differ among PFGs.

The 3D configuration of cells in a landscape affects each of the microenvironmental parameters – for example, a high-rise building casts shade on neighboring cells, and gaps in the terrain can prevent the spread of fire. The model makes no assumptions regarding the spatial configuration of cells (with the exception of seed dispersal and neighboring biomass, see below), but expects that all inputs follow the same pattern. It further maps the outputs back to the same landscape representation.

There are two further processes that occur on the landscape level. First, newly produced seeds are dispersed to other cells. Due to the intended small spatial extent of the landscape, it is assumed that seeds are redistributed uniformly across the whole site, possibly aided by zoochory. The 3D configuration of the landscape thus has no bearing on seed dispersal. Secondly, one experimental setting allows light to fall at an angle, and pass through neighboring cells. In this case the spatial configuration has a direct influence on competition outcomes.

In the stand-alone version of the model, all cell attributes are constant throughout the model run, i.e., there is a fixed annual disturbance in each cell, and the habitat suitability is only determined at initialization. However, the model is also compiled as shared library, allowing it to be embedded in other programs and models (not available under Windows). If using the ECOLOPES PLANT MODEL as part of a different program, the inputs “soil class” and “disturbance” can be exchanged at every time step. This allows, e.g., modelling directional and stochastic climate change through changes in the soil class attribute.

#### 3. Process overview and scheduling

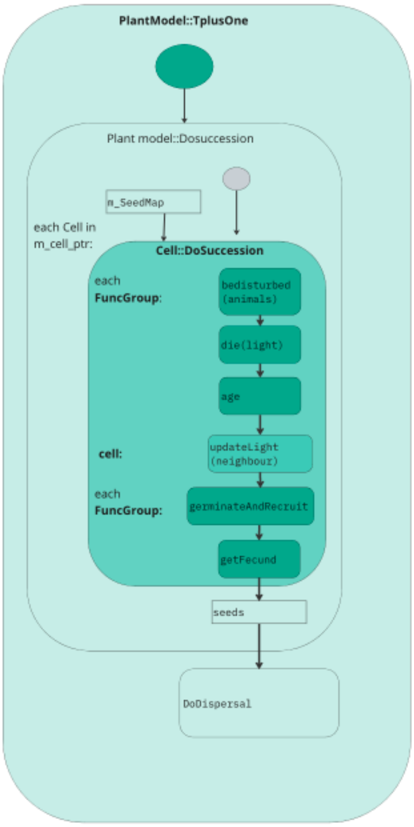


*Fig. 1: Process overview. The order of events is largely copied from Fate-HD, so that conceptual changes to the model are kept to a minimum. Processes in this diagram match function names of code (see UML diagram).*

The model largely works on individual cells, and reflects the process that is also employed by Fate-HD (Fig. 1): In each time step, each deme (FuncGroup in the code) is disturbed by a number of fixed, annual disturbances. Each PFG may react differently to the disturbances, including not being affected at all. Then it is checked, separately for each stratum, whether the current light conditions are sufficient for the FuncGroup. A common outcome is the death of all lower-growing cohorts (indicating lower plant parts), while the upper cohorts survive (see also issue #21). Subsequently the deme ages by one time step, potentially causing the oldest demographic group to die. Aging is followed by a calculation of the new light conditions. Light is calculated based on the demography of the whole community (not a single FuncGroup) of the cell, and in contrast to FATE-HD, also of the neighboring cell if shading by neighbors is enabled. Germination and recruitment are then individually computed for each FuncGroup, and the outcome is a number of newly produced seeds. These processes depend on the current suitability of the cell, which is in turn determined by soil depth and soil class (but see issue #22). The newly dispersed seeds are not immediately placed back into the cell. Instead, all seeds across the landscape are collected and then uniformly and randomly dispersed across the site.

### 4. Design concepts

#### 4.1 Basic principles

FATE-HD (Boulangeat et al., 2014), from which the ECOLOPES PLANT MODEL is derived, combines phenomenological and process-based approaches: it first considers abiotic environmental filters to determine where each agent (deme) can in principle live, and then simulates their growth and demographic structure, including competition among agents. The ECOLOPES PLANT MODEL adds flexibility to the phenomenological approach, and allows users to derive habitat suitabilities instead based on other (process-based) models. It further lets the processes be influenced by explicit (as opposed to conceptional and generalized) inputs, such that future versions can accurately predict biomass in absolute terms.

In preparation for accurate predictions of biomass, light is conceptually treated differently. “Light competition” and “shading” in Fate-HD in fact represent integrated (and unitless) canopy effects. In this model “shading” is taken literally, allowing to calculate light that traverses neighboring cells and to include shading by geometric structures such as buildings. The implications of this conceptual change are further below (see “calculation of current light resources”, section 7.2).

#### 4.2 Emergence and interaction

The key outcome of the simulation is the spatiotemporal change of community composition, which emerges from the interaction of demes of different PFGs within each cell and stratum (particularly competition for light). On a landscape scale, abiotic factors (soil depth, soil class) impose a general community structure via habitat filtering, and these abiotic factors may be static (inputs) or dynamically changing (passed on from another model). The abiotic factors (shading, in particular) further affect some competition outcomes, leading to the emergence of complex environment-community relationships, but I consider these indirect effects secondary and weaker than those imposed by habitat filtering.

#### 4.3 Adaptation

The plant model does not contain any direct objective-seeking mechanisms, i.e. the agents do not possess the ability to choose among two or more alternative behaviors e.g. via learning or condition-dependent rules (phenotypic plasticity).

#### 4.4 Objectives

The plant model does not simulate adaptive behavior (fitness objectives). The rules by which agents respond to the environment (the "genotypes" of the Plant Functional Groups) are instead provided by the user.

#### 4.5 Interaction

Currently, individual agents (demes) do not interact with one another, directly or indirectly, except in the competition for light, which is explained in detail in the submodel section (see also issue #23). Briefly, growth and death of age cohorts continually changes light conditions, and, subject to light tolerances, cohorts may be outcompeted and die. This leads to turnover of plant communities.

#### 4.6 Stochasticity

The ECOLOPES PLANT MODEL contains two sources of stochasticity:

1. At the end of the time step, all plant seeds are pooled and randomly redistributed over the simulation environment, together with the seeds that enter from the region (seed rain). This simplified form of dispersal was chosen over the pre-existing disc-based dispersal from FATE-HD, because the spatial extent of the site is expected to be smaller than the usual dispersal kernel of all PFGs.

2. The initialization seeds a random number of the age cohorts 0 and 1 years to each cell where the habitat is generally suitable. The cohort size follows a random uniform distribution between 0 and 100.

Habitat suitability is a stochastic process in FATE-HD (cells have a certain probability to be suitable), but in this version of the ECOLOPES PLANT MODEL I opted for purely deterministic behavior (see issue #48). Stochastic behavior can still be provoked by adding a (separately compiled) stochastic soil class model that changes the soil class in each time step; this allows for much more flexibility and setting of own rules, e.g. gradual change, autocorrelated biased random walks, etc.

#### 4.7 Implementation

FATE-HD is an agent-based model, but not implemented in traditional format. In agent-based models, simple rules are set up for individual entities (agents), and the interaction among agents causes complex behaviors to emerge. In ecology the agents are usually individual cells or individuals of a population, which may for example grow, compete, reproduce and die. Individual agents are nested in habitats, populations or environments (multiple levels of nesting may occur), and the nesting is usually reflected in a Object-Oriented Programming (OOP) approach. Moreover, individual instances of the lowest level are created (birth) or deleted (death), and may be reassigned to other instances of higher-order objects (dispersal). The modular structure of the OOP approach allows modification of agent rules easily.

In FATE-HD, on the other hand, the principal agents are age cohorts, i.e. groups of individuals with the same age. The individuals hence are not uniquely identifiable, and there is no inter-individual variation or plasticity in individual decisions; essentially this moves the modelled entity to the population (or deme) level. The deme (vector of age cohorts) is nested within a plant community of a cell, and the community/cell is nested in an environment. Because only the age cohorts are created, deleted or changed, there is a single instance of each deme within each cell, which is constructed at initialization and only deleted at program termination. This approach is computationally more feasible, but comes at the cost of model resolution and accuracy (only the community composition, but not individual plant development can be tracked). This model inherits the concepts, including treatment of individuals, from FATE-HD, but revised the model architecture to allow easier inclusion of novel structures later.

##### 4.8 Software engineering approaches

FATE-HD follows field-specific software engineering principles of ecological models. See (Scheller et al., 2010; Vedder et al., 2021) for an overview of common approaches, potential drawbacks and risks. The ECOLOPES PLANT MODEL, on the other hand, is specifically designed as a platform that is planned to be further extended in the future. This prototype is likely to be used in applied settings (e.g., architectural practice, urban planning) and embedded in a complex workflow. The current version is thus neither accurate nor neccessarily correct (see section 9), but built to be consistent, maintainable and extensible. The model thus employs more robust software engineering principles than usually found in ecological modelling, including the following measures:

1) the model architecture was revised to achieve stronger encapsulation and modularization. In particular, classes no longer access data that should be outside of their scope;
2) the code was modernized to include c++17 and c++20 features, thus achieving better readability, higher maintainability and code safety; removal of GDAL and boost libraries further ensures portability to other platforms;
3) tests, assertions and unit tests ensure that the model works as intended; the tests are embedded in a CI system that also includes up-to-date documentation of the code with doxygen;
4) the inputs, output and internal data formats were changed to allow handling of 3-dimensional data; all files are in standard JSON format;
5) the model can be compiled as library (on linux and MacOS), allowing combination with other C++ models. Habitat suitability (soil) and disturbance information can be exchanged by other models at runtime.

Conversion of the code is not finished, however (see issues #25-#31).

#### 5. Input and Output Data

The ECOLOPES PLANT MODEL requires the following information to run:

- Configuration parameters (Table 3)
- PFG attributes, including their response to disturbances (e.g. management, animals) and soil (soil classes, depth requirements) (Table 4)
- Microenvironmental information for each cell. This currently includes the amount of shading, soil class and soil depth, and the amount of each (user-defined) disturbance.

The output of the model is the biomass of each PFG in each cell, for each time step that was specified. A list of time steps to output can be provided via the configuration parameters.

See appendix Supp. S2 for details on input and output data formats and their use.

*Table 3: Configuration parameters. N indicates the number of parameters that need to be supplied.*

| Name | N | Description |
|---|---|---|
| SimulDuration | 1 | Number of time steps to simulate |
| SaveYears | - | List of time steps for which output is saved |
| NoStrata | 1 | Number of height strata (usually 4) |
| StrataHeight | - | Describes height of each stratum |
| PotentialFecundity | 1 | Multiplication factor for fecundity (table 2) |
| MaxAbundLow / Medium /High | 3 | Maximum abundance small/medium/large PFGs can reach |
| SeedingInput | 1 | Number of additional seeds introduced every time step |
| LightThreshLow / Medium | 2 | Abundance threshold upon which light conditions decrease |
| LightAngle | 1 | Angle of the sun in degrees (range 0-90) |

*Table 4: PFG parameters. N indicates the number of parameters that need to be supplied.*

| Attribute | N | Description |
|---|---|---|
| Maturation time | 1 | Years from birth to fecundity |
| Life span | 1 | Years from birth to death |
| Maximum Abundance | 1 | Abundance class (small/medium/high) of this PFG |
| Immature size | 1 | Proportion of immature plants relative to mature abundance |
| Max stratum | 1 | Which stratum(layer) does the mature plant reach? |
| Stratum change age | 1-3 | At which ages does the plant reach the next stratum(layer)? |
| Seed pool life span | 2 | Life span of seed pools (seeds decrease exponentially) |
| Potential fecundity | 1 | Rejuvenation potential of a PFG. |
| Shading factor | 1 | How much shade does one unit of biomass produce |
| Germination probability | 3 | Max germination probability under low/medium/high light conditions |
| Light tolerance | 9 | Survival of each life stage under each light condition.<br>Life stages are:<br>Propagule(ignored), germinant, immature, mature<br>Light conditions are: low, medium, high |
| Soil tolerance | - | List of soil classes on which PFG can survive |
| Soil depth requirement | 1 | Minimum soil depth in cm that is required for PFG survival |
| Disturbance response | - | Effect of each disturbance on mature and immature plants, respectively (percentage biomass destroyed). Read via separate file |

### 6. Initialization

Upon startup, the model reads all inputs, and then reserves memory and creates all demes in all cells. First, the model reads the input file, which contains the file name of the global simulation parameters(“GlobSimulFile”), and the file name of the data file (“DATA”). The model then reads and checks the information in those two files as well. Using the information in the global simulation parameters and in the DATA file, the model checks the data content, and builds and initializes the cells and their FuncGroups.

Building the demes includes seeding a random number of plants (aged 0-1 years) to each cell where the habitat is generally suitable. The number of plants per age group follows a random uniform distribution between 0 and 100.

The plant model runs 5 iterations during initialization. Running the model for several iterations ensures a buildup of a reasonable plant community, as the cells are initially seeded with only 0-1 year old plant material. An option to input a more detailed starting population is currently missing (issue #18).

### 7. Submodels

The modelling approach contains three main processes: habitat suitability, succession, (including disturbance, death, aging, germination and recruitment and reproduction) and dispersal.

#### 7.1 Habitat suitability

The habitat suitability module of Fate-HD checks for each PFG in which cells it can theoretically occur (habitat filtering), based on a static user input. The ECOLOPES PLANT MODEL instead uses a soil class that can be provided as a static input by the user or as a dynamically changing property provided by a different model. The soil class of each cell is compared with the soil class requirements of each PFG (each PFG can potentially live on multiple soil classes), and it also checks whether the soil depth is deeper than the minimum requirement of each PFG. The outcome of this comparison indicates the suitability of the cell for each PFG.
Although treated as soil depth and soil class throughout the model and its description, the calculation is based on simple integer comparison and list comparison, respectively. It can easily be repurposed for other means, by providing a different meaningful integer or string input, and by changing the PFG attributes accordingly. For example, one may input the groundwater table as “soil depth”, and a PFG’s root length as “depth requirement” - The model then does the sensible comparison whether root length is larger than groundwater level. One may similarly repurpose the “soil class” and call the inputs “contaminated soil” and “clean soil”, “low humidity”/”medium humidity”/”high humidity”, or any other factor one deems important for suitability. The PFG attributes must then be changed accordingly.
The suitability of a habitat impacts whether seeds can germinate. Already established plants will not be destroyed if a habitat becomes unsuitable (issue #22).

#### 7.2 Succession

Succession describes the growth processes that happen within a year inside a single cell. It consists (in this order) of disturbance, death due to insufficient light, aging, calculation of current light resources, germination and recruitment, and reproduction (Fig. 1). While the implementation of succession was revised, the general calculations and concepts have not changed (yet). Long-term seed dormancy is disabled, however (see issue #26 for an explanation).

##### Disturbance

The disturbance model of FATE-HD allows to provide a map (for one disturbance) or several maps (several disturbances) for each PFG and each stage (juvenile, mature) that can either cause a given % of mortality or a given % of seed production. The lost material may, however, grow back at the end of the time step, depending on the type of disturbance.

In the ECOLOPES PLANT MODEL, I make two kinds of disturbance possible:
1. the user may provide a constant disturbance map that works largely the same way as in the original version, and
2. the user may provide a new map every time step. The dynamic maps are not read from files, but passed on from another model as c++-internal objects. This requires incorporating the plant model library into another project.

In the PFG definitions one can define the effect each kind of disturbance has on each PFG. The constant maps represent management decisions such as annual tree pruning, while the dynamic maps are, for example, annually changing herbivore biomasses. Currently only one of the two disturbances can be used at a time.

In contrast to FATE-HD, the idea of resprouting and of seed mortality were both given up, i.e. a disturbance can only remove an amount of abundance/biomass/canopy.

1. Seeds in FATE-HD are not meant to be an accurate integer representation of produced seeds, but rather comparable to a general “seeding” or rejuvenation function; removing a particular percentage of a general function did not seem reasonable to parametrize, and the seed pool is anyway in a state that warrants rewriting (issue #26).
2. Resprouting essentially causes some part of the plant canopy to become temporarily invisible (for the light calculation) and then reappear at the end of the time step. Rewriting of the succession order would likely break resprouting, so it was not yet added to the model. Furthermore, resprouting appears to confuse demographic and individual-level processes: a number of individuals is removed from the population demography, but a biomass regrows towards the end of the time step (see also issue #29 and section “succession”).

##### Death due to insufficient light

This function works on the light conditions of the preceding year. Light is expressed as a vector of factors, classifying the light in each stratum as low, intermediate or high. It is tested for each cohort whether the light tolerance matches the current light conditions, given the current growth stage (seed, germinant, juveniles, matures). The 12 light tolerance parameters, i.e., low, med and high for each growth stage, are provided in the PFG definitions. Cohorts that do not tolerate the current light conditions die.

The code defines cohorts as a group of individuals with the same abundance but differing in age (i.e., the exact opposite of a cohort in the usual sense). In order to work on age cohorts as described above, it requires splitting the "Cohort" (c++ object) into actual age cohorts. The process is further implemented as recursive function through changing an iterator of a loop, which adds to the complexity (Fig. 2), is computationally expensive and error-prone (see also issue #25).

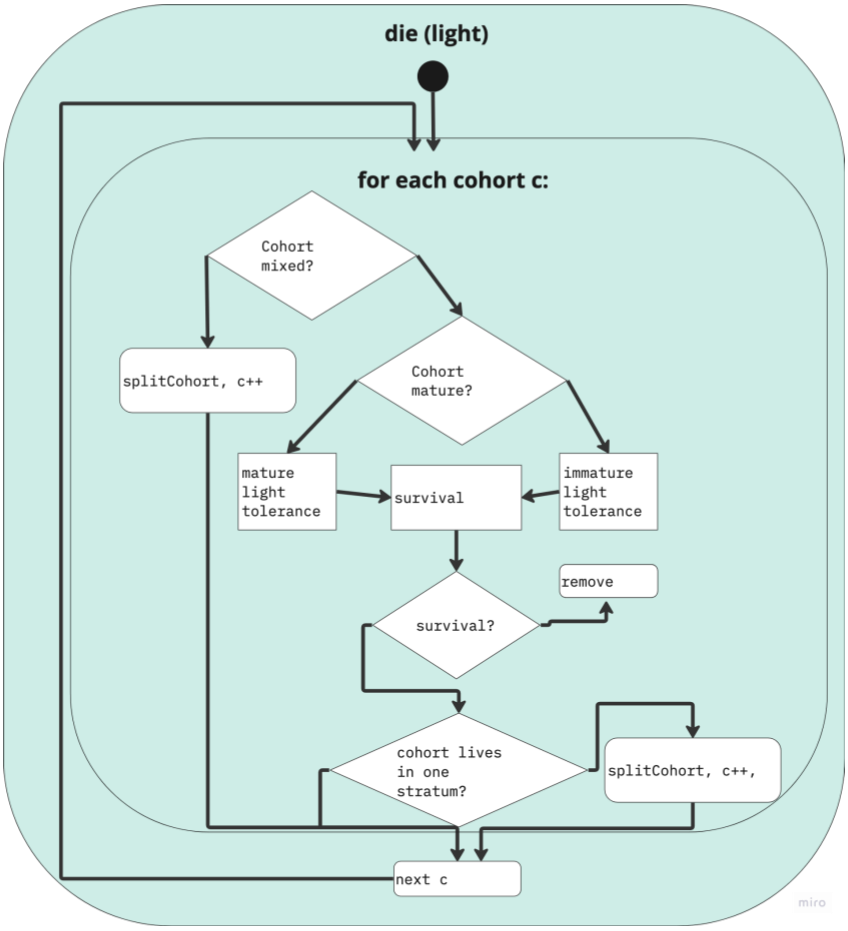


*Fig. 2. Description of death by insufficient light. Note that "cohort" in the code does not refer to a biological age cohort.*

##### Aging

The description of the demography of the functional group is updated, according to the LifeSpan and maturity attributes of the PFG. Material that exceeds the life span of a PFG is removed.

#### Calculation of current light resources

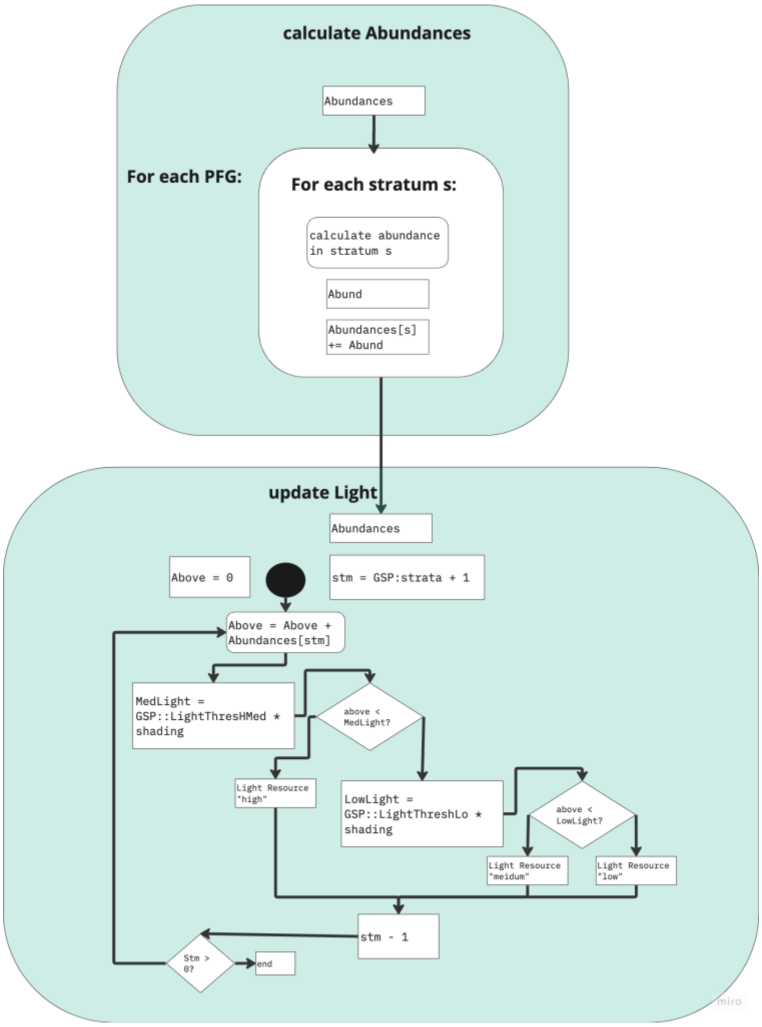


*Fig. 3: calculation of light resources.*

The technical implementation of this submodel mostly follows the logic of FATE-HD:
The light is calculated for the uppermost stratum according to the biomass of the cohorts living in it. The light that is not consumed is passed on to the next stratum, being reduced in the same fashion until it reaches zero or arrives at ground level (stratum 0). Consumption of light depends on a shading factor (not to be confused with the input "shading") that represents leaf and growth form, maturity of the cohort, and on the biomass that the PFG reaches (the maximum biomass that a plant can reach differs between small, medium and large plants, and the size class is another PFG attribute). The amount of light in each stratum is converted into a factor (low, medium and high light conditions). In terms of technical implementation, the model does not use light, but a ray of „shading" passes through the cell. It collects any (leaf-form and maturity- corrected) abundance it encounters. In each level the shading ray is compared against a light threshold parameter, and if smaller than the "low" or "medium" threshold, the light resource declines to the according level.

Although called light/shadeFactor in FATE-HD, the collected abundance ("shading" above) is meant to represent canopy effects, i.e., the integrated effect of shading, humidity changes, and other abiotic and biotic factors. In the ECOLOPES PLANT MODEL light/shading was taken

literally, and hence has a slightly different meaning. For instance, in Fate-HD it was perfectly logical if an understory plant required a canopy for survival; in the ECOLOPES PLANT MODEL it would not make biological sense if a plant could survive under shaded, resource-limited conditions, but would die under full-light conditions that contain abundant light resources. In other words, the survival under shade but not under full light would be exclusively a competition outcome in the ECOLOPES PLANT MODEL, but the result of habitat filtering and facilitation in FATE-HD.
Using the more literal interpretation of light/shading, the ECOLOPES PLANT MODEL was able to make two further additions to the concepts and implementation.
First, and in contrast to the original Fate-HD implementation, the threshold against which the shading ray is compared is corrected by a shading index (a proportion ranging from 0 = full sun to 1 = full shade). This ensures that shaded cells fall earlier to "medium" or "low" light conditions. Due to the multiplicative nature of the calculation, the uppermost stratum can never be converted into lower light levels (multiplication of zero shade with a proportion). This could theoretically cause issues in the calculation of light in the uppermost stratum in very shaded positions (see issue #20); however, plants are always expected to tolerate full light anyway (see above).
Secondly, light may fall at an angle below 90°. If this is the case, if will partially pass through the neighboring cell and is accordingly reduced by the neighbor's plant abundance. The calculation is as following:

Light will enter into stratum i, get depleted by the plant biomass, and the remaining light enters stratum i-1 etc. (Fig. 4A). When light enters at an angle (Fig. 4B), a proportion bypasses the upper layer. Knowing the height of the stratum (a) and the angle at which light enters(ß), we can calculate the length at which light hits the ground as x = a/tan(ß) (x is actually an area, not a line, but if light enters exactly from south/east/etc., the calculation does not change). With a cell size of 1m, the area not covered by light is y= 1-x. The light falling on the ground is thus y * (light from above) + x * (light from above in neighboring cell). We can repeat this calculation for every layer. The angle ß is a simulation parameter that can be changed, but the direction of incoming light is fixed at  x  = - inf.*

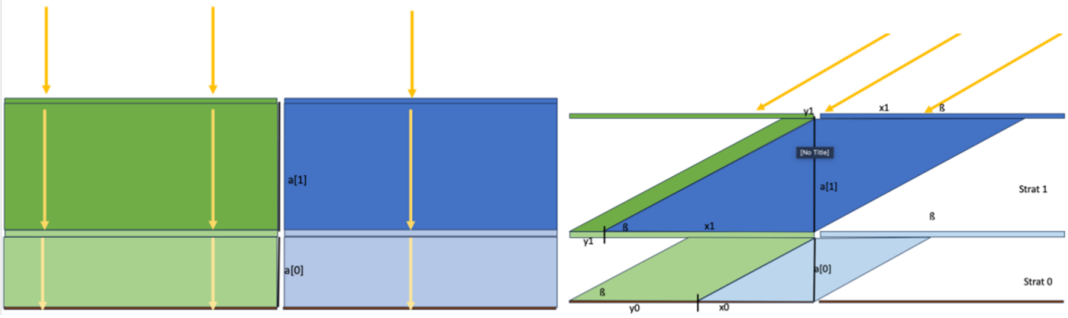


*Fig. 4. Light calculations of Fate-HD (A) and new plant model with angles activated(B). A: Light passing through voxel cells as in current model version and in Fate-HD. B: Methodology for light passing in at an angle. The proportion of light falling into stratum 0 but coming from the neighboring cell is x1=a[1]/tan(ß), and the proportion of light going through this cell's upper layers is y1 = 1-x1. The calculation is repeated for every stratum.*

#### Germination and recruitment

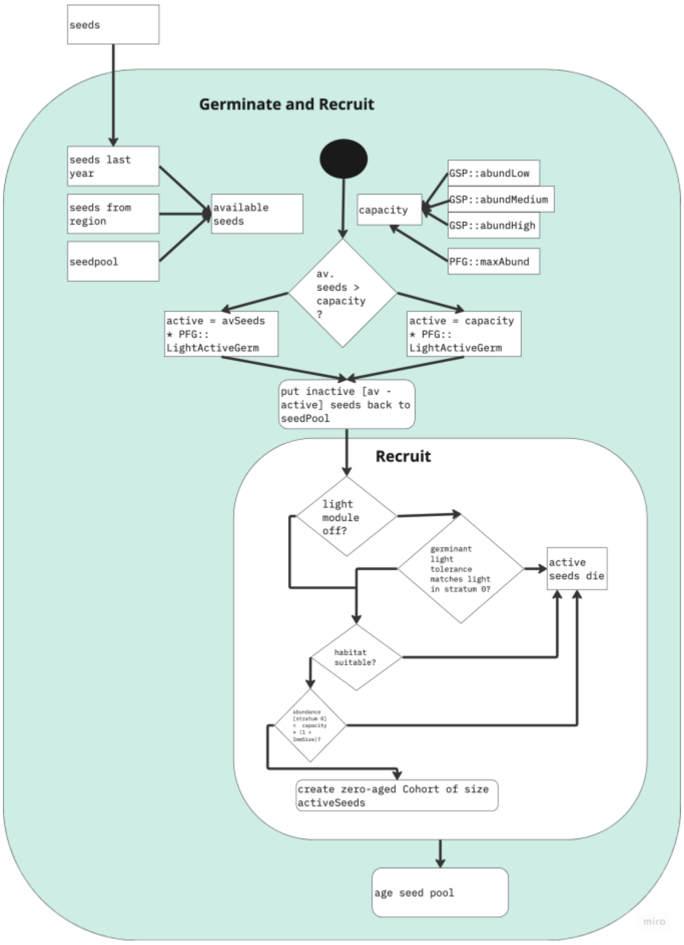


*Fig. 5: Germination and recruitment.*

The seeds from which new plant biomass is recruited consists of 1) last year's seeds that landed in the cell; 2) seeds from the existing seed pool (buried in the soil); and 3) seeds arriving from the region. The number of new seeds is checked against the soil's capacity for each deme, which is a configuration setting. If the number of seeds is lower than the current capacity, all seeds are activated, otherwise the excess seed mass is put back into the seed pool. All activated seeds then enter the recruitment function, and when light conditions are sufficient, habitat is suitable and there is capacity left, zero-aged plant material is produced. Any activated seeds for whom the above conditions are not met die and are permanently removed from the model.

#### Reproduction

The reproduction function returns either the current abundance of mature cohorts, or the cell's capacity (whichever is smaller), but scales the value with a fecundity parameter (a configuration setting). If fecundity is left at 1, the seed abundance produced in each time step is as large as the current (adult) deme biomass. This ensures massive overproduction of seeds, and plants will regrow whenever conditions are suitable.

##### 7.3 Dispersal

In FATE-HD the dispersal model was based on three parameters: d50 that is the maximum distance within which 50% of the seeds are dispersed, d99 is the maximum distance within which 99% of the seeds are found in total, and ldd is the maximum long-distance dispersal. Within each dispersal disc, the seeds could be distributed uniformly or decreasing exponentially with distance from the deme.
Plant dispersal has been simplified in the ECOLOPES PLANT MODEL in comparison to FATE-HD: all seeds produced in a year are pooled, and then redistributed randomly across the site at the end of the year.

#### 8. Implementation

| Concept | Detail |
| --- | --- |
| Operating system | Portable to any common OS |
| Programming language | c++ 20 (or newer) |
| Precursors | FATE-HD; some code (data containers, inputs) shared with joint ECOLOPES models |

Being developed as part of a joint model and partially derived and altered from FATE-HD, the model contains code and concepts that are currently not required and better turned off (e.g. seed dormancy), and code which works but uses outdated programming paradigms. Special care is taken that the documentation is up to date, but some code comments or documentation sections may ocassionally be forgotten.

##### 8.1 Compiling as library or executable

The model can be compiled as a library (.dll or .dylib), so that it can easily be #include'd into another program. This requires copying the library file and the header file "plantmodel.h" to the target directory (or its "include" folder) and linking against it. The root directory contains a “mockup.cpp” which provides a demonstration. See the .yaml file for compile instructions that do not require CMake. In addition to using as library, one can compile the source files together with main.cpp (in src folder) to create a full executable.

There is furthermore a "tests" subfolder, in which unit tests according to the GoogleTest framework are conducted. The test procedure is also included in the CMake files.

Because this is work in progress, compiling is untested and may not work yet under Windows (issues #6 and #16).

##### 8.2 Dependencies and requirements

All dependencies are included in the /include folder (ECOLOPES joint model for data containers, nlohmann::json and easylogging). The model has no specific requirements to run (< 1 GB hard disc space, < 2GB RAM, any modern CPU), but compilation requires CMake and a c++20 compiler (it is tested on llvm and MinGW's GNU compiler, but should also work on AppleClang). See Supp. S2 for installation notes.

#### 8.3 Implementation overview

The ECOLOPES PLANT MODEL consists of individual Cell objects (10,000 for a 100x100 site). Each cell contains abiotic information (soil class, shading), as well as a plant community that consists of multiple FuncGroups (one per PFG). The FuncGroups store the demographic information of the PFG in the cell. The ECOLOPES PLANT MODEL is also linked to various inputs, consisting chiefly of 1) general simulation parameters (GSP), 2) constant user inputs (e.g. shading, soil depth), and 3) constant PFG definition (containing e.g. Life span). The input classes are inherited from a base class, which all submodels of the ECOLOPES ecological model should share.

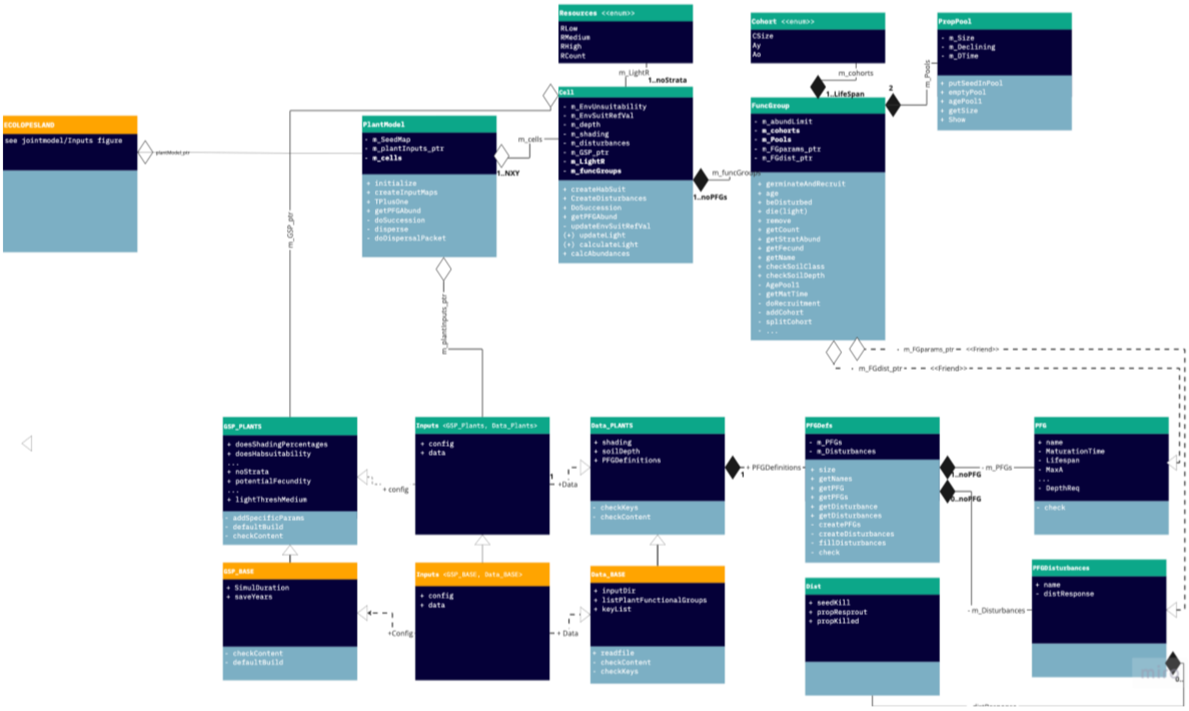


*Fig. 6: UML diagram of plant model. The upper half contains the model itself, the lower half classes for inputs (including PFG data). The input classes of this model as well as for separate animal and soil models are inherited from a common parent class (orange). Some smaller enums are not included to avoid cluttering. Notation follows standard UML notation. "EcolopesLand" refers to a class of a different model, which may #include the plant model as a library and run it.*

### 9 Warnings and limitations

The model is not fully parametrized and not validated. Inputs are assumed to be coarse approximations or simplifcations, and the model accordingly only suggests relative differences among demes. The model is not yet thoroughly tested and currently meant to be used only for academic purposes.

#### 9.1 Problems, bugs, inconsistencies

This is an incomplete list of the most important conceptual and technical issues. Please refer to the issues section on GitLab (https://gitlab.com/ecolopes-team/plantmodel/-/issues) for an up-to-date and complete list. Various of the issues listed here will be solved with a new individual-based model (issue #29).

##### Shading issues (issues #20, #21, #23)

The original plant model was built for a landscape-scale. The community of each cell was a stand of trees etc, containing multiple individuals that may shade each other. By reducing the spatial extent to a square/cube meter, we now may have only a single indivdidual of a PFG present. The highest stratum of the PFG shades the lower parts and causes them to die, which is no longer realistic when referring to a single plant.
Additionally, plants that grow within the same stratum do not compete with each other for light, only plants reaching a higher stratum will compete with others.

##### Other conceptual issues (issues #22)

When the soil changes, it does not affect growing or mature plants, it only affects the seed recruitment process. One could also let it affect the fecundity of the plants. In the original RFate implementation this was possibly handled differently. If a habitat turned unsuitable, the FG would be treated like an immature, so the fecundity would be set to zero. It seems as if it was also treated in the light calculation like an immature (small) plant, yet it still aged normally. This might be a conceptional oversight of the original implementation.

##### Cohorts and legions (issue #25)

The model simulates age cohorts and does not include inter-individual variation. This was a reasonable choice made to simplify the model complexity. This section does not deal with the conceptional choice per se, but about the way in which the cohorts are implemented.
A simple way to store the demography is to use a vector of size "lifespan" to indicate the abundance of each age group. For example,

[ 10, 10, 10, 20 , 20 ]

Would indicate 10 plants each aged 0-2, and 20 plants each aged 3-4. FATE-HD (and hence also the ECOLOPES PLANT MODEL) instead chooses to summarize the information to (0, 2, 10) and (3, 4, 20), i.e., a vector of cohorts with the information (minAge, maxAge, amount). I see various limitations of the approach:

1) This variable does not contain the information of one age cohort, but information across age cohorts. It is the exact opposite of what its name conveys and it should be renamed.
2) This way of storing information might in principle save a bit of memory, but I doubt that the amount is large (or even positive). In the example with 5 age classes, the cohort notation requires more memory (six integers). If a cell contains 3 strata, it is likely that there will be at least three cohorts, which are differently affected by the light conditions. This makes 3*3 = 9 integer values to store, so the approach can only save memory if the median life span of the PFGs is at least 10. If the maturity age does not coincide with the strata, or there are more strata to model, the number of integers to store increases further.
3) While the classical implementation (1 value per life span) may be more expensive when a plant does not occur (many zeroes), it has the benefit that it has to be constructed only once per cell and PFG. The need for manual memory management (a cumbersome topic in c++) would be completely avoided, and one would not need to worry about memory leaks or memory fragmentation.
4) The chosen way of storing and dealing with data requires rather complex operations whenever a cohort passes over an age border. For example, a tree may reach its mature size at age 10. In year 9, there is a cohort of (0, 9, 20); in year 10 the cohort needs to age and become (1, 10, 20). Then an offspring cohort (0, 0, 20) is created and subsequently joined (0, 10, 20). If a disturbance is applied to immature plants only, the cohort is split at age 10 into (0, 9, 20) and (10, 10, 20). In year 11 all of those steps are repeated, but the split cohort (10, 10, 10) is added to the new (11, 11, 10) cohort. All the time the model deals with a vector of cohorts (size 1-2 in this example). Resizing a vector or inserting something into the middle of a vector is prohibitively costly and increases computation time. More importantly, the code needs to be maintained, tested and debugged, which increases the development time.
5) The way cohorts are split and put together is also suboptimal in my view. A for loop/while loop is used to loop over the cohorts, and whenever a cohort is split, the iterator needs to be adjusted. This results in a recursive-like function (Fig. 2). It is customary that one should not change loop iterators, and although it is technically possible it can easily result in logical errors that are hard to debug.
6) The model code would in principle also allow for overlapping cohorts; for example, one could have 50 individuals each from age 0-10; 20 individuals aged 3-5, and 40 individuals aged 4-7. This notation is equivalent to 50 individuals aged 0-2; 70 individuals aged 3; 110 individuals aged 4-5; 90 individuals aged 5-7; and 50 individuals aged 7-10. This freedom to choose different cohorts for the same demography was possibly not intended, and may affect the calculations (it does seem to affect the age() function).

To make the code more maintainable (and enhance performance), the cohorts are disassembled at each time step ((0,0,10), (1,1,10) etc.) and reassembled at the end, but the system should be entirely revised at some point. The disassembly has the side effect of removing cohort overlap.

#### PropPool (issue #26)

The seed propagule pool is not well described in either the original paper or the user manual, so it is a bit difficult to infer the concepts and ideas behind it. Essentially it seems like a functionality that has long been abandoned. As I understand it, the amount of seeds produced every year depends on the abundance and fecundity of a PFG. Only a certain proportion of the seeds can germinate every year, so naturally a seed pool would build up in the soil (there is no carrying capacity or density dependence in the seed pool). This is counteracted by a constant seed mortality rate which diminishes the seed pool over time. However, if the user chooses a long "shelf life" of the seeds, one would still build up a considerable seed bank, and because recruitment is proportional to the seed bank, a considerable rejuvenation potential (potentially a conceptual oversight?).

The implementation of the PropPool class is a bit awkward to use and raised a few questions. The class consists of a seed set ("m_size") that is meant to decline with a constant mortality rate, and this mortality rate is given by

size (n+1) = size (n) - size(n) * (1 / (pl + 1)),

where pl is is the pool life span. The decline happens whenever the member function "agepool1" is called. For instance, with a pool life span of 1 the seed set halves every time step, with a life span of 10 years it decreases by 10% per year. This means that the seeds diminish exponentially.

The pool life span is not a property of the class, but a PFG attribute, passed on as function argument whenever the pool ages. This means that the responsibility for the pool life span has been shifted away from the object and to the caller (who may e.g., decrease the seed pool by 50% in year 1, but by 10 % in year two), which is suboptimal.

The class also contains a private "DTime" variable, which is never used, and the declining of the seed set can be disabled (bool m_declining) which does not seem to make much sense. The class contains a public "emptyPool()" function which erases all content but does not properly delete the object.

Lastly, the class contains a "putseedsinPool" function, which replaces the seed set by a new one, if the new one is larger than the older (it does not add the seeds but replaces them). If the new seed set is smaller than the old one, the function does nothing. I do not understand why this approach was chosen. The age of the seeds is not tracked (or rather not used?), and mortality is independent of seed age anyway (it depends on life span, not age), so one could simply have added the seeds. I think this might be a bug. This bug seems to counteract the lack of a carrying capacity, as the seed pool is emptied and refilled (but only to a maximum value) every time step.

I suggest rewriting and making the mortality rate a const argument that cannot be changed after instantiating the object. The constructor should implement the linear equation based on pool life span, so this function is only called once per seed pool object. I would remove the

m_declining variable, and also the emptyPool() function. The class FuncGroup should instead delete the pool and replace it by a different one if required. One could alternatively create a “addSeedsToPool” function.
It also seems that the function FuncGroup::agepool1(), which indirects to PropPool:: agepool1() is not being used. I think it should be erased.

That said, seeds are somehow stored, and if not germinating they will age and die. Only seed dormancy is better left disabled.

##### Enums and Fracts (issue #27)

The plant model contains various unscoped enumerations (enums), such as Abund (can contain the values “low”, “medium” or “high”) or LifeStage (“Propagule”, “Germinant”, “Immature”, “Mature”). These enums are internally stored as integers and can be implicitly converted (can be verified by hovering over an enum in vscode). Standard enums are neither type safe nor strongly scoped, so “Abund::low == LifeStage::Propagule” would be true but not sensible (or even “Mature – Immature == medium”). It is better to use scoped enumerations (enum **class**).

The plant model further uses enums to circumvent calculations based on floats. In general, calculations based on integers used to be faster than calculations on floats (“Fract” enums). Because Enums are internally stored as integers, the float essentially is rounded to an integer. These enums have 10 levels (10, 20... 100) and their use reduces the precision but may in principle increase performance. I see two problems with the approach: 1) there is no performance gain between an enum of size 10 and an enum of size 100, so one could round to the nearest integer instead of the nearest decade; 2) CPUs are no longer limited by processing time but mostly by loading the memory. If speed is an issue, one may simply convert from double to single precision, which is still several orders more accurate than the fract conversion.

Multiple functions are required to convert between doubles and Fract’s, bloating the code and requiring additional function calls. I suggest removing the fracts in the future.

##### Vector-map conversion (issue #28)

The model used to run on std::vectors. In light of the intended use (long-term development and maintenance), we opted for the slightly slower but safer std::maps approach. Most parts have been changed to std::maps to reduce the possibility of confusing inputs/assigning attributes to the wrong plant functional group, but some remnants of the old vector system still exists, and any conversion risks breaking something. Proper tests with multiple PFGs are required (tedious), or, even better, the remaining vectors of the SuFate class need to be cleaned up.

#### 9.2 Ideas for the future

##### Light calculation and 3D in the plant model (issue #17, #15, #29, #31)

The plant model is, strictly speaking, only correct when all cells are horizontal and light enters at a 90° angle. This is unrealistic, in reality light comes in at a variety of angles (changing over the course of the day), and the distribution of angles is latitude dependent. When the cell is tilted, the distribution of angles is strongly affected by the aspect (north/east/south/west) of the cell. When light enters at an angle, it does not pass through all strata of a cell, but partly through strata of a neighboring cell. A first version of light passing through the immediate neighbor is implemented, but limited to a single light angle rather than a distribution, and limited to horizontal and immediately adjacent cells.

The above already allows for better 3D effects, causing e.g. vegetation on a building edge to receive more light than vegetation in the center. There are a few disadvantages though: 1) if light comes in at a sufficiently steep angle, it should pass through multiple cells. This is ignored in above version; 2) with this method one can only model light coming from the eastern/southern etc. neighbor, but not light coming in from, e.g., SSW. This is important when trying to simulate a natural distribution of light angles; 3) the technique does not work with sloped angles or even facades, as tilting 3D cells will cause them to overlap and make calculations impossible.
A solution to above problem is raytracing, I.e. following the sun rays and checking with which cells they collide. This technique is employed in any modern 3D video game ("shaders/shading") but potentially would slow down the model considerably.

The addition of shading through neighbors made it necessary to implement a cell height variable. There are no checks yet that two stacked 3D cells do not overlap (issue #31)

##### Solving the shading issues (issue #29)

Most of the conceptual and technical issues highlighted above (competition with self, cohorts) relate to the lack of plant individuals in the model. A new model version could contain a plant community that is made up of individuals; the individual's biomasses (distributed across strata) convert collected light into a resource and store it in a pool (root system); the resource is used for maintenance as well as to create new biomass and new seeds. Thus, shading on the lower parts of the plant are compensated by the higher parts of the plant. Another major change in the new model is replacing any factors by integers. In the new model light, measured in lux, is converted into biomass, measured in kg. This allows correctly parametrizing the model with real-world data.
