## Supplementary material for "The ECOLOPES PLANT MODEL : a high-resolution model to simulate plant community dynamics in cities and other human-dominated and managed environments": Supp. S2

### The ECOLOPES Plant Model – user guide

Jens Joschinski, Studio Animal-Aided Design

This report has been written by Jens Joschinski. Some text is copied or adapted from (non-public or public) documents of the ECOLOPES project. ECOLOPES has granted permission to use the text presented herein.

Copyright (C) 2022 - present Studio Animal-Aided Design
Permission is granted to copy, distribute and/or modify this document under the terms of the GNU Free Documentation License, Version 1.3 or any later version published by the Free Software Foundation; with no Invariant Sections, no Front-Cover Texts, and no Back-Cover Texts. A copy of the license is included in the section entitled "GNU Free Documentation License".

This software has been developed as part of the ECOLOPES project.

#### Installation notes and settings

The plant model requires at least c++ 20 to compile. It is guaranteed to compile on clang, and MinGW's GNU compiler for windows ("gcc"). The program is best compiled with CMake, a CMakelists.txt is available (building with cmake . -G "MinGW Makefiles"  should help under windows). The repository on Gitlab contains compiled windows and linux executables: go to the Menu CI/CD->pipelines to find a download button for an up-to-date version.

If building from source, check the (potentially hidden) file ".gitlab-ci.yml" for commands on linux and win.

Full command line instructions to install all programs and compile on MacOS:

Brew upgrade

Brew install cmake

Brew install llvm

cd path/to/work_directory

cmake -B build -DCMAKE_CXX_COMPILER=/usr/local/cellar/llvm/16.0.4/bin/clang++  <--insert correct llvm version here (check in Finder)

cmake --build build

cd out

./Model_Darwin_1.0.1

Regardless of the OS, CMake will create a new directory "/out", in which all model and test executables can be found. In addition, a dynamic library is built (.dylib, .so or .dll). With the help of the public header in include/plants/, the library can be #included in other programs.

#### Running

To run, the model requires a json file with file names/paths to find the inputs, and with some global simulation parameters.
By default, the work directory is root/tests, where root is the work directory in which the model was built. The default filename is test.json.

To change default paths and file names, use:

model.exe -w path/to/dir -f filename

(see model.exe -v for more command line options)

The subfolder /tests already contains a full example, including a test.json. The json files can be opened with any text editor, but opening with VSCode or Mozilla Firefox ensures that the files are displayed in a pretty format.

The test.json contains all neccessary data (order is not important):

- the simulation duration (5 iterations)
- which iterations shall be saved (only the results from years 1,2 and 4 are saved)
- input and output directories ("IN" and "RESULTS") relative to the working directory
- listPlantFunctionalGroups belongs to a partially implemented feature, which allows selecting only locally available PFGs. Currently this list should contain exactly the same information, alphabetically sorted, as the PFGDefsFile (see below)
- the name of a "mask" file from which the coordinates or cell names are copied (this is currently an independent file, mask.json, but could also be set to e.g. management.json)
- the name of a file that determines what kinds of management occur where on the site (management.json)
- the microenvironemntal parameters, soil class, shading and soil depth. The soil file is straightforward, open ninesoil.json to have a look. You will find nine coordinate-triplets, each with a soil class associated. The shading and depth files are coded differently in this example. Go to tests/nine.json to check the file content. There are also 9 coordinate keys in the file, but each key is associated with two values (called SHADING and DEPTH). Hence, the entry in test.json has two fields per microenvironmental variable: a file name (nine.json in both cases) and a field name ("SHADING" or "DEPTH").
- the filename of a logger configuration file. No need to change unless you know what you are doing.
- The last two important entries in test.json are the PFG and disturbance definitions. the PFG definitions describe what kinds of PFGs can theoretically occur on site, and defines their traits. The disturbance file describes for each PFG, which management plans or other disturbances (e.g. herbivory) may affect the growth of the plant. Not all disturbances need to occur on a site, and not all disturbances that occur on a site need to affect a PFG. Hence it is difficult to check the inputs for errors automatically. Please take special care regarding the spelling of disturbances, and mind capitalization.
- further global parameters, such as the number of strata and max size of the plants.

Having read the input information, we know that the model will run with 2 PFGs and for 5 iterations, on a 3x3x2 ecolope. We now hit

```
model_win_1.0.1.exe -f test.json
```

The model will finish within one second, but relevant information has been printed on the terminal (and in separate log files). This includes the reading and content of major files, warnings encountered while trying to read, and information regarding the time steps. The model saves various information while running. The results can be explored unter RESULTS, which is the output directory that was written into test.json. Each of these files contains 18 values, corresponding to the 18 cells of the site.

Due to the input, the following things will happen in the cells:

- in various cells there is not enough soil to support trees (0,0,0, 1,2,0 etc.)
- in cell 0,0,0 there is 1.0 fire. Because grass is heavily affected by fire (100% killed), no grass will grow there. In all other cells, there is 0.5 fire, causing biomass of grass to decline by 50% per time step
- in cells starting with a 2 there is 100% shading, so grasses that are not shade-tolerant cannot grow (2,0,0 etc). Trees are shade-tolerant and can grow if there is enough soil.
- cell 2,2,1 does not have a suitable soil for either plant
- in cells that do get tree growth, light is insufficient for grass due to light competition

So let us look at the results (assuming the RNG seed is fixed):

- cell 0,0,0 has no trees (soil depth) and there is 100% fire. However, seed pressure from neighboring cells seems to be high enough to get 5000 plants per year. This number is the maximum abundance a plant of class "highAbund" (PFGDefs)such as grass can attain (see test.json::maxAbund)
- cells 0,0,1 and 0,1,0 can support trees (enough soil depth), causing grass to decline to 0. The amount is up to 5000 per stratum (highAbund Plant as well) + up to 5000 seeds (20000 total)
- cell 0,1,1 has no trees and fire is reduced in comparison to 0,0,0. Here grass can grow nearly twice as high (up to 9542).
- all cells with a leading 2 have either no plants at all (insufficient depth for trees) or only trees, due to the shading
- cell 2,2,1 is an exception, as it would support trees in terms of soil depth, but has the wrong soil class (hence also no growth at all)
